# Integrating learning in movement using step-selection analyses

**DOI:** 10.64898/2026.09.01.747820

**Authors:** Benedetta Catitti, John R. Fieberg, Martin U. Grüebler

## Abstract

1. Understanding how animals acquire and use information from the environment is critical for linking movement to population dynamics, species distributions and conservation. Advances in tracking technologies and growing interest in learning processes have opened opportunities to study behaviours such as habitat exploration in translocated animals or ontogeny of migration and dispersal movements. However, accessible statistical methods for studying these behavioural processes are lacking.
2. We present a learning-explicit step-selection analysis (SSA) that integrates movement, habitat selection, and learning into a single evolving process. At the start of the movement trajectory, the animal is assumed to have no knowledge of the landscape, i.e. its internal habitat quality map is initialized to a constant. As the animal moves, this map is updated dynamically: with each step, only the habitat within its perceptual range becomes known and contributes to future decisions.
3. Through simulations, we show that this method separates true habitat preferences from learning effects and reveals when large-scale behaviours, such as attraction to resources or avoidance of risks, emerge as knowledge accumulates. Critically, we demonstrate that ignoring learning, as in traditional SSA, can lead to biased estimators of habitat selection.
4. Finally, we apply our approach to a real-world GPS dataset of naïve individuals, consisting of 10 juvenile red kites dispersing in Switzerland. Using leave-one-individual-out cross-validation, we show that learner SSFs consistently outperform traditional SSFs in predicting movement decisions.
5. We conclude by discussing methodological considerations and future directions for integrating learning into movement ecology. By explicitly modelling the formation of memory that integrates both spatial information and habitat quality, our approach advances process-based movement ecology while remaining compatible with standard SSA workflows, offering practical tools for both theoretical research and applied wildlife management.

## 1 Introduction

Understanding how animals move in their environment is crucial for uncovering the mechanisms underlying species distributions and population dynamics, and for designing and implementing effective conservation strategies (Abrahms et al., 2021; Allen & Singh, 2016). This information need, along with advances in animal biotelemetry, have fuelled the development of new statistical methods and software for modelling the environmental drivers of animal movement (Joo et al., 2022). More recently, considerable attention has been given to the role of learning and memory in driving animal movements, reflecting that animals base their decisions not only on intrinsic factors (e.g., age, sex) or external conditions (e.g., habitat quality) but also on knowledge gained from past experiences. The influence of past information has been explored using mathematical and statistical models to explain the observed spatial patterns of site fidelity in foraging, breeding, and use of refuge areas (Bracis et al., 2015; Gurarie et al., 2022; Iorio-Merlo et al., 2022; Schlägel & Lewis, 2014). Although these approaches focus on how stored information (i.e. *memory* in the form of “cognitive maps”) is used, process-based approaches that model how information is collected (i.e., *learned*) are scarce, primarily for two reasons. First, most empirical movement data have been collected on adults, due to technical and economic concerns linked with tagging juveniles (e.g. reduced natural survival). When using data from adults, it is challenging to model the learning process without knowledge of the individual’s past experiences (i.e., prior to when the animal was tagged). Second, mechanistic models that include learning typically require the addition of spatiotemporal parameters as well as customised likelihood functions—making implementation challenging for most ecological research settings (Falcón-Cortés et al., 2021; Thompson, Derocher, et al., 2022; Thompson, Lewis, et al., 2022). Recent advances in high-resolution GPS technology, and concurrent reductions in device prices, have enabled detailed investigations into juvenile movement behaviour and natal dispersal dynamics (Catitti et al., 2024; Debeffe et al., 2013). Effective conservation translocations also depend on understanding how relocated individuals acquire and utilize spatial information in novel environments (Falcón-Cortés et al., 2021; Frair et al., 2007). Thus, the development of accessible analytical frameworks to quantify learning processes would provide valuable insights for both theoretical ecology and applied wildlife management.

Step-selection analysis (SSA) has emerged as one of the most widely used frameworks for studying animal movement from tracking data, offering a powerful integration of movement mechanics and habitat selection. At its core, SSA decomposes movement into discrete steps by combining a movement kernel, which characterizes movement capacity through probability distributions of step lengths and turn angles, with a selection function that quantifies habitat preferences at each potential destination (Thurfjell et al., 2014). This dual structure enables researchers to disentangle movement constraints from active habitat choices, providing mechanistic insights into space-use patterns (Fortin et al., 2005; Thurfjell et al., 2014). The method’s popularity stems from its flexibility in modelling spatiotemporal dynamics of habitat use, its accessibility through standard statistical software implementing conditional logistic regression (Signer et al., 2019), and its active development by the scientific community (Fieberg et al., 2021; Hofmann et al., 2024; Klappstein et al., 2024; Michelot et al., 2024; Signer et al., 2024).

Recent developments aimed at integrating memory processes in SSA have focused primarily on adding “familiarity functions” that capture an animal’s tendency to revisit previously explored areas (reviewed in Kim et al., 2024). These functions typically include metrics such as the intensity of use of past locations, angular deviation from prior movement trajectories, or the time elapsed since the last visit to a location. However, these methods are designed for adult animals with prior spatial knowledge, requiring the exclusion of an initial portion of the tracking data as a “burn-in” period to account for unknown pre-tracking experience, along with assumptions about the appropriate length of this period (Avgar et al., 2015). In contrast, when using data of juvenile animals that start naïve to their environment, no such pre-existing spatial memory needs to be accounted for, allowing the full trajectory to be used for analysis and the actual learning process, and the resulting build-up of memory, to be investigated.

A second limitation of current SSA-based approaches is that they often focus on fine-scale, local movement decisions at the step level without adequately capturing how learning influences broader-scale movement patterns. Because step-selection functions operate in discrete time steps and evaluate choices against a local set of available steps, they inherently limit inference to local motivations. To bridge to larger-scale movement drivers, existing approaches typically add specific attraction or repulsion terms toward known features or areas, formulated using distance-to or angular covariates, which again presumes some form of learned past spatial knowledge (Prokopenko et al., 2017). In contrast, juvenile animals provide a unique opportunity to investigate when large-scale behavioural shifts occur as a result of learning, such as a sudden change in directionality of movement following the discovery of a valuable resource. Failing to account for the broad-scale implications of the learning process risks misattributing or biasing habitat-selection effects. For naïve or translocated animals, modelling movement as if an important landscape feature were always known can underestimate its true biological influence, not because attraction to the feature is weak, but because its effect is mediated by learning over time. Critically, this underestimation propagates to the broad scale: movement patterns that arise from delayed awareness and information acquisition may be poorly captured when models assume constant knowledge, leading to mismatches between observed and predicted large-scale movement paths.

We propose a streamlined analytical framework that integrates a movement kernel, habitat selection, and learning within a SSA framework to model the behaviour of naïve individuals. Rather than adding complexity, our approach simplifies the model structure by merging habitat selection and learning into a single, evolving process. At the start of the movement trajectory, the animal is assumed to have no knowledge of the landscape, i.e. its internal habitat quality map is initialized to the mean value of the landscape. As the animal moves, this map is updated dynamically: with each step, only the habitat within its perceptual range becomes known and contributes to future decisions. In a first step, we use this generative framework to simulate trajectories of naïve, learning individuals in two scenarios: one in which animals move through a heterogeneous habitat and select for high-quality areas, and another that additionally includes a discrete attraction source (an “Easter egg”; Fig. 1, step 1). The latter scenario allows us to examine how large-scale movement patterns emerge following the discovery of localized resources and to show that such patterns can be understood correctly only by accounting for the fact that resources must first be encountered before they can influence movement decisions. We then compare a traditional (“omniscient”) SSA, which assumes complete knowledge of the environment, with a learning-explicit (“learner”) SSA fitted to the simulated trajectories. This comparison reveals potential biases arising from the omniscience assumption and evaluates whether they can be reduced by explicitly modelling learning (Fig. 1, step 2). Finally, we apply our method to empirical data of naïve individuals from juvenile red kites *Milvus milvus* (Fig. 1, step 3). We then discuss the ecological contexts in which this learning-explicit approach offers insight, along with associated methodological considerations and avenues for future research.

**Figure 1.**
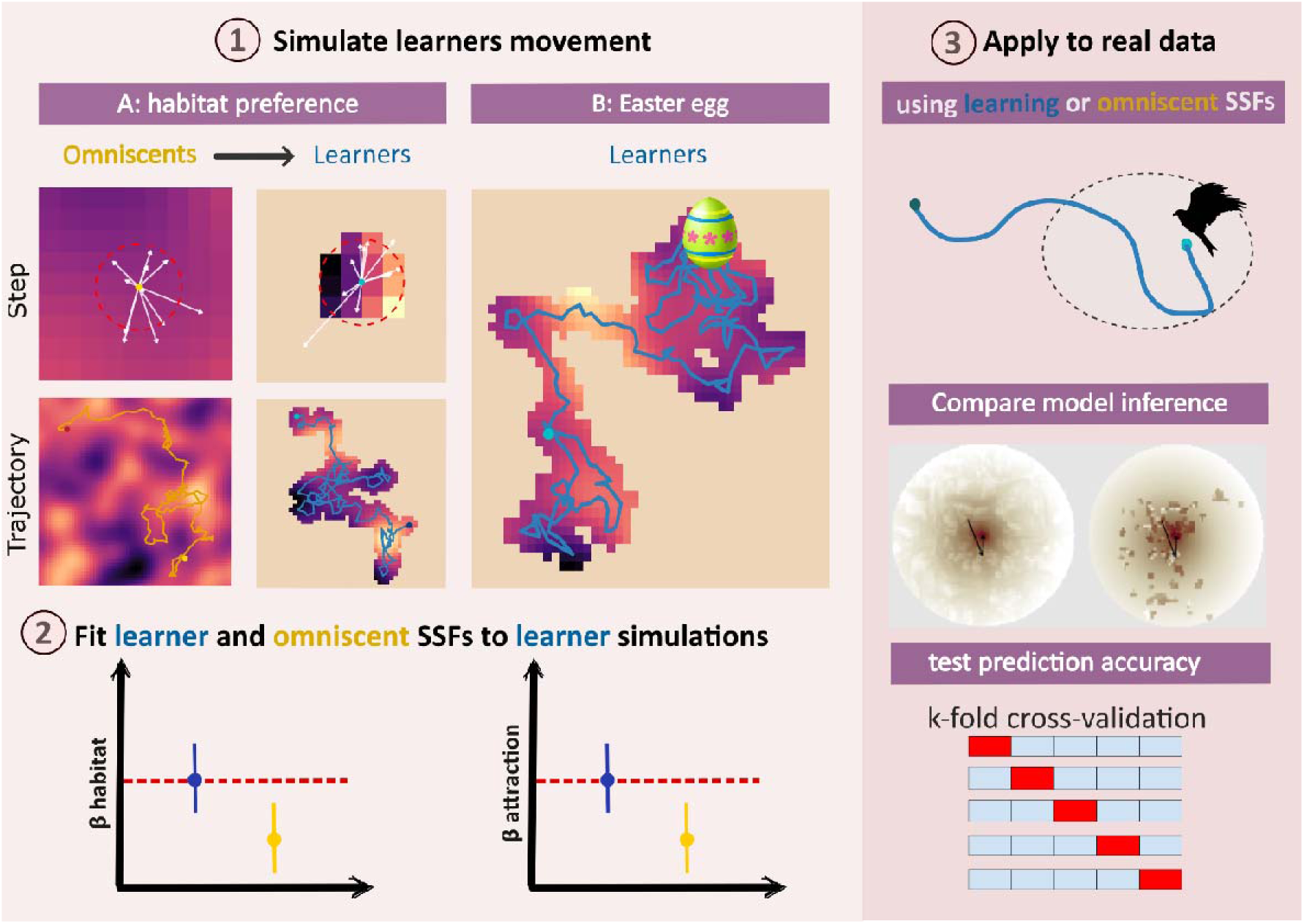
We explore the importance of incorporating learning in SSAs using a variety of analyses. First, we simulate 500 movement trajectories for both “omniscient” and “learning” individuals, with positive selection for an underlying habitat covariate (scenario A). Omniscient individuals follow a traditional approach and have full knowledge of the environment from the outset, whereas learner individuals are initially naïve and progressively acquire information as they move. We extend this framework to a scenario B that also includes an attractive source (an “Easter egg”) in the landscape (scenario B). Next, we then fit SSFs to the learner-generated trajectories to test whether selection coefficients are more accurately recovered by models that explicitly incorporate learning compared to traditional omniscient SSFs. Finally, we apply both SSF approaches to empirical movement data of 10 dispersing juvenile red kites. Model performance is assessed using AIC and leave-one-individual-out cross-validation by comparing predictive accuracy between learner and omniscient formulations.

## 2 Simulation of learner and omniscient tracks

### 2.1 Methods

#### 2.1.1 Simulations

We simulated 500 movement trajectories representing a *naïve* individual (hereafter “learner”), such as a juvenile dispersing from its natal home range or a translocated animal encountering an unfamiliar landscape. We simulated movement in two habitat scenarios (Fig. 1A-B). In both scenarios, individuals selected for a spatially heterogeneous habitat covariate. In scenario B, we also added a discrete resource to the landscape capable of eliciting extreme attraction, such as a key resource patch, hereafter referred to as an “Easter egg”.

In both cases, i.e. (A) the habitat-only and (B) the habitat + Easter egg scenarios, movement and habitat –selection parameters were defined *a priori*. We specified a positive selection coefficient for favourable habitat values (β habitat = 0.5). To represent limited environmental knowledge typical of naïve individuals, the habitat layer was initialized to its global mean value (0). The true habitat values were revealed only within a fixed (here set to 1 unit) perceptual radius around the individual’s current location, reflecting the assumption that the individual initially perceives only its immediate surroundings and progressively learns about the environment as it moves (Fig. 1). In the habitat + Easter egg scenario, we additionally specified a positive coefficient (β = 0.8) associated with the relative angle to the Easter egg. We used angular deviation rather than distance to because it provides greater contrast among candidate steps within a choice set and more directly represents directed movement toward a focal target. Each trajectory was initiated from a fixed starting location with a randomly assigned initial orientation. At each movement step, we generated a set of 100 candidate steps by drawing turn angles from a uniform distribution and step lengths from a gamma distribution. For each candidate step, we extracted the value of the perceived habitat at the endpoint. When a candidate step terminated in an area not yet revealed, a habitat value equal to the global mean (0) was assigned, reflecting the absence of prior knowledge about that region. Additionally, in the habitat + Easter egg scenario, the response to the Easter egg was modelled as state dependent: individuals were assumed to be initially unaware of the egg, and the corresponding selection term was activated only after the individual’s perceptual range intersected a 7-unit buffer around the egg, which defined the detection zone. The radius of this buffer can be interpreted as a feature-specific detection distance and may vary according to the extent or conspicuousness of the source (e.g. a larger buffer may be appropriate for a noisy water source, such as a stream, versus a less detectable pond).

For each candidate movement step, we computed standard step-selection covariates, including step length (sl), log-transformed step length (log(sl)), and the cosine of the turn angle (cos(ta)), which facilitate estimation of movement parameters associated with gamma and von-Mises distributions (Avgar et al., 2016; Fieberg et al., 2021; Michelot et al., 2024). Using the predefined selection coefficients, we calculated a relative selection score *w*(x) for each candidate step. These scores were normalized to obtain selection probabilities, from which a single step was probabilistically selected and executed. After each realized step, the habitat layer was updated within the perceptual radius centred on the new location, thereby expanding the individual’s known environment (Fig. 1). This procedure was iterated for 2,000 steps per trajectory, yielding movement paths that emerge from the interaction between intrinsic movement tendencies, habitat selection, and spatially explicit learning of environmental conditions over time. Trajectories were constrained to reach 2,000 steps. When a trajectory reached the boundary of the environmental layer, the simulation forced the selection of a new step whose heading remained within the layer.

#### 2.1.2 Comparison of movement characteristics for learner and omniscient trajectories

To test whether this learning process leads to movement characteristics that differ from those of individuals assumed to have perfect knowledge of their environment, we additionally generated an equivalent number (i.e. 500) of simulated trajectories for both scenarios using a traditional mover with perfect knowledge of the landscape (hereafter “omniscient”) and otherwise identical movement parameters.

We compared basic movement metrics, including 100% Minium Convex Polygons (MCPs), revisitation rates and residence times, associated with learner and omniscient trajectories simulated under the habitat-only scenario. MCPs were calculated using the *st_convex_hull* function in the *sf* package (Pebesma, 2018). Revisitation and residency patterns were quantified using a recursion-based approach via the *recurse* package (Bracis et al., 2018), whereby circular areas of increasing radius (1–5 units) were centred on each location along a trajectory, and subsequent returns to these areas were recorded. For each simulated track and chosen radius, we calculated the mean number of revisits and the mean residence time within each circle. Lastly, we compared the tortuosity of the learning-based and omniscient simulations. To do so, we quantified local path tortuosity along each simulated trajectory using a sliding-window approach. Tortuosity was defined as the ratio between the total path length within a segment and the straight-line distance between its start and end points, such that higher values indicate more convoluted movement. For each trajectory, tortuosity was calculated over overlapping segments of fixed length (window size = 100 consecutive locations). Tortuosity values were computed for all valid segments along a track and then averaged to obtain a single mean tortuosity estimate per simulated individual. MCPs, recursion patterns and tortuosity were summarised using means and 95% Confidence intervals (95% CI).

We evaluated these same metrics in an additional set of simulations (500 learner and 500 omniscient individuals) conducted in a more homogeneous habitat (Fig. S1). These simulations allowed us to assess whether learner and omniscient individuals differ in their exploration of the environment, and to determine whether any such differences depend on the degree of environmental patchiness.

### 2.2 Results

Simulated movement trajectories differed between learners and omniscients in overall space use and movement characteristics (Fig. 2). Learners tended to occupy slightly larger areas, although differences were modest (learner mean MCP = 7469 units^2^, 95% CI = 7288–7651; omniscient mean MCP = 7113 units^2^, 95% CI = 6940–7286; Fig. 2A–B).

**Figure 2.**
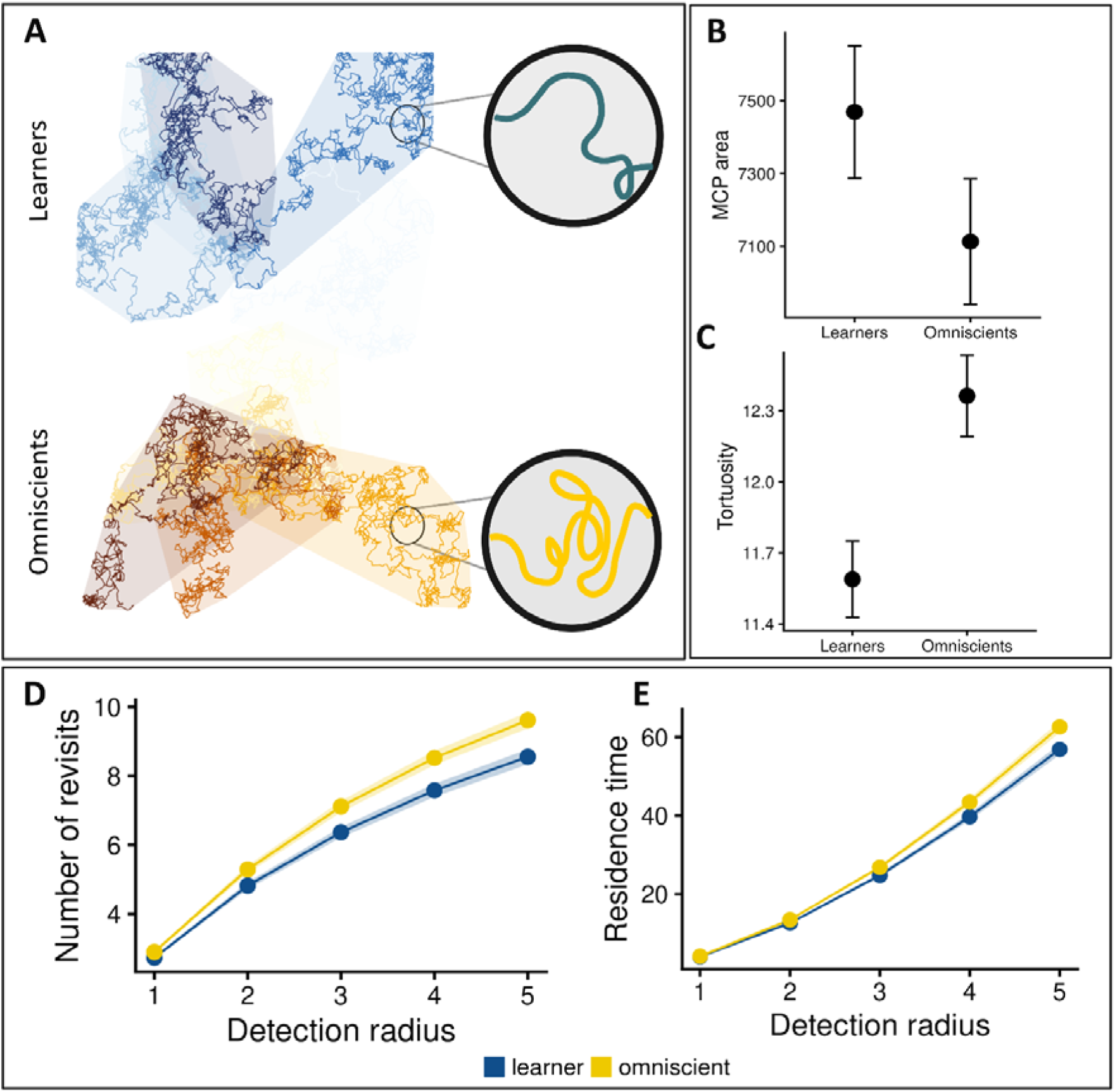
Comparison of movement metrics derived from 500 simulated trajectories under omniscient and learner movement strategies. Movement trajectories were simulated with a positive selection for favourable habitat values (β habitat = 0.5). In the traditional, “omniscient” scenario, individuals select their next step by evaluating environmental conditions for all available steps. In the “learner” scenario, individuals have only partial knowledge of the environment: candidate steps are either known (previously visited or within perceptual range) or unknown. (A) Example of five simulated trajectories for each movement strategy. (B–C) Comparison of space use and movement structure between strategies, quantified as 100% minimum convex polygon (MCP) area (B) and path tortuosity calculated over a moving window of 100 consecutive steps (C). Differences between groups were assessed using a non-parametric permutation test. (D–E) Comparison of movement recursiveness across increasing perceptual radii, quantified as the number of revisits (D) and residence time (E)

Mean path tortuosity was higher for omniscient individuals than learners, indicating more sinuous movement paths (Fig. 2C). The difference between the two groups was modest with high certainty (learner mean tortuosity =11.6, 95% CI = 11.4–11.7; omniscient mean tortuosity = 12.2, 95% CI =12–12.5; Fig. 2C).

Revisitation rates increased with the size of the radius used to define a revisit (for both learners and omniscients), but were consistently higher for omniscients. For learners, the mean number of revisits increased from 2.73 (95% CI = 2.68–2.77) when using a radius of 1 unit to 8.55 (95% CI = 8.34–8.77) when using a radius of 5 units, whereas for omniscients the mean number of revisits ranged from 2.90 (95% CI = 2.85–2.94) to 9.61 (95% CI = 9.38–9.85) for those same radii (Fig. 2D). Mean residence time within revisited areas showed a similar pattern, increasing from 4.03 (95% CI = 3.97–4.09) to 56.9 (95% CI = 55.5–58.2) for learners and from 4.17 (95% CI = 4.11–4.23) to 62.6 (95% CI = 61.2–64) for omniscients (Fig. 2E). Thus, omniscients revisited locations slightly more frequently and remained within revisited areas longer than learners. Differences between learner and omniscient trajectories were less pronounced in a more homogeneous habitat. As the patchiness of the environment decreased, there were little to no differences in tortuosity and residence time between the two groups (Fig. S1).

## 3 Application of SSA to learner trajectories

### 3.1 Methods

We fitted step-selection functions using both the traditional (omniscient) approach and our learner approach to the 500 learner-simulated tracks in the habitat-only and habitat + Easter egg scenarios. With the learner approach, the habitat layer is dynamically updated, replacing mean values with known habitat covariates within the perceptual range of the individual as it moves, and the attraction towards the Easter egg only starts upon its discovery. In the omniscient approach, the individual is assumed to know the habitat and the Easter egg at all locations. We quantified bias in both approaches by comparing the mean estimate to the “true” habitat-selection coefficients (β_habitat_= 0.5; β_angle_ = 0.8). We expected differences between learner and omniscient models would be largest during the initial phases of movement, when naïve animals have the least amount of information. Towards the end of the trajectory, we expected the two approaches to result in unbiased, or nearly unbiased, estimators. To test these predictions, we used *mgcv* and *gratia* packages (Pedersen et al., 2019; Simpson, 2024) to fit step-selection functions with time-varying habitat-selection parameters (Klappstein et al., 2024). Specifically, we fitted a step-selection function adding as covariates step length, the log of step length, the cosine of turn angle as typical SSF movement kernel covariates (Fieberg et al., 2021) and a smoother interaction between the step id (our time covariate) and habitat (habitat-only scenario), and additionally a smoother interaction between step id and the relative angle to the Easter egg (habitat + Easter egg scenario). To fit step-selection functions in the latter scenario, we only retained trajectories where the Easter egg was detected at least once (N = 218). As a comparison, we also fitted non-time-varying step-selection functions to each of the 500 simulated learner tracks using the *amt* package (Signer et al., 2019). In the habitat-only scenario, we additionally evaluated the learner step-selection function across a number of perceptual ranges, from smaller (0.25, 0.5, and 0.75) to larger (1.25, 1.5, 1.75, and 2) than the value used to generate the trajectories. We then used AIC to select the best performing model among the omniscient and learner models with various perceptual ranges (Akaike, 1974). This allowed us to evaluate the effect of the broad modelling approach (omniscient vs learners across different perceptual ranges) on habitat-selection coefficient estimators and determine if the perceptual range could be identified using standard model selection procedures.

### 3.2 Results

#### 3.2.1 Habitat-only scenario

When fitting a traditional (omniscient) step-selection function (SSF) to the simulated learner trajectories, we consistently underestimated the underlying habitat-selection coefficient (Fig. S2; β̂ = 0.285, 95% Monte Carlo interval [0.276, 0.294], compared to the true value β̂ = 0.5). Furthermore, in our time-varying coefficient model, we observed a gradual increase in the estimated habitat-selection coefficient over time, from β̂ = 0.206 [0.184, 0.228] at step 5 to β̂ = 0.340 [0.317, 0.363] at step 1955. As expected, the strongest underestimation in the time-varying model occurred at the beginning of the trajectory. At this stage, the animal has limited knowledge of its environment; however, a traditional SSF, assuming full environmental awareness, interprets these initial movements as reflecting weak habitat selection rather than incomplete information. As the animal acquires knowledge through movement, the estimated coefficients progressively increase, but over 2000 steps they do not converge to the true habitat-selection coefficient.

In contrast, a learning-based SSF accounts for the animal’s information constraints. At each step, the observed location is compared to 100 random alternatives, for which habitat values are known only if they fall within the animal’s perceptual range or have been previously visited. Because this framework mirrors the data-generating process used in the simulations, it recovers the true coefficient (β̂ = 0.499 [0.495, 0.504]) and yields stable estimates over time in the time-varying model, from β̂ = 0.492 [0.481, 0.504] at step 5 to β̂ = 0.506 [0.494, 0.519] at step 1955 (Fig. S2).

The learner step-selection function recovered the habitat selection coefficient most accurately when the assumed perceptual range matched the true one (1.0; Fig. S3). The learner model assuming a perceptual range of 1.0 had the lowest AIC for 498 out of 500 simulated individuals (Fig. S3). Support declined monotonically as the assumed range departed from the true value in either direction. The omniscient model (mean ΔAIC 89) was outperformed by every learner model except the one with the smallest perceptual range (mean ΔAIC 99).

#### 3.2.2 Habitat + Easter egg scenario

When fitted to trajectories generated under the learning-explicit process, the omniscient SSF again underestimated both components of selection: the habitat-selection coefficient and the coefficient describing directed movement toward the Easter egg (Fig. S4). This bias was evident in models that estimated a single constant coefficient across the full trajectory (habitat β̂ = 0.317, 95% Monte Carlo interval [0.299, 0.334], true value 0.5; source-directed movement: β̂ = 0.616 [0.591, 0.640], true value 0.8), as well as in models with time-varying coefficients (habitat: from β̂= 0.222 [0.184, 0.259] at step 5 to β̂ = 0.457 [0.422, 0.491] at step 1955; source-directed movement: from β̂ = 0.181 [0.136, 0.227] to β̂ = 0.778 [0.762, 0.794]). As in the habitat-only scenario, the early portion of the trajectory showed the strongest discrepancy between the fitted omniscient model and the underlying data-generating process, consistent with the fact that individuals were initially unaware of habitat quality and the Easter egg, but notably improved as the trajectory progressed (Fig. S4). Coefficients were well recovered by the learner model, both in its constant form (habitat: ββ̂= 0.501 [0.490, 0.511]; angle to the Easter egg: β̂ = 0.793 [0.786, 0.801]) and its time-varying β form (habitat: β̂ = 0.502 [0.480, 0.523] at step 5 and β̂ = 0.488 [0.461, 0.516] at step 1955; angle: β̂ = 1.08 [0.52, 1.64] at step 5 and β̂ = 0.798 [0.782, 0.814] at step 1955). In the time-varying coefficient model, uncertainty associated with the selection for the relative angle to the Easter egg was large early in the trajectory and declined thereafter, reflecting heterogeneity in the timing of detection across simulations and the corresponding delay in activation of the source-response term.

## 4 Application to real-world data

### 4.1 Methods

We compared the two step-selection approaches (omniscient and learner) using data from 10 juvenile red kites during their first dispersal event from their natal areas in Switzerland (i.e., from the moment they leave their parental home ranges until their first migrations). We first restricted analyses to daytime locations (7AM–7PM local time) and resampled the tracks to an approximately regular 1-hour schedule (with 15 minutes tolerance). We assumed that, in the absence of habitat selection, step lengths followed a gamma distribution and turn angles followed a von Mises distribution. To estimate parameters, we formed strata by matching each observed step to 100 randomly generated steps using empirically fitted gamma and von Mises distributions. We then fit a conditional logistic regression model with step length, log step length, and cosine of turn angle and two habitat covariates (slope and NDVI). Slope (60-m resolution) was derived from a DEM-based raster (https://opentopography.org/) and NDVI (30-m resolution) was obtained from Sentinel-2 L2A imagery using the *rstac* package (Simoes et al., 2021). We centred and scaled both rasters so that they had a mean of 0 and standard deviation of 1 prior to covariate extraction at step endpoints. We compared two information assumptions: an omniscient SSF in which environmental covariates are assumed to be known everywhere (static raster values), and a learner SSF (new method) in which environmental information is locally available and updated through time within a specified perceptual radius around the animal’s current position, producing time-varying “known” covariates. We evaluated multiple perceptual radii (10, 50, 100, 250 and 500 m) spanning a range from well below to somewhat above the median observed step length. We kept the interval broad in order to test whether AIC and cross-validation can identify an appropriate perceptual range in the realistic case where the true value is unknown.

We compared the omniscient model to the learner models with different perceptual ranges using two complementary criteria. First, as in the simulation, we ranked models by AIC. Second, we computed a leave-one-individual-out cross-validated log-likelihood (i.e. an out-of-sample log score): for each bird, we summed the log-likelihood contributions of its strata, evaluated using coefficients estimated from the other nine birds. For each held-out individual and perceptual range, we then took the difference in summed log likelihoods between the learner and omniscient models, so that positive values indicate better prediction by the learner SSF.

### 4.2 Results

Individuals selected for higher NDVI values, with the estimated strength of selection dependent on whether individuals were assumed to be omniscient or learners, and in the case of learners, dependent on their assumed perceptual range (Fig. 3). These results are consistent with those obtained from our simulation study, where analysing the movements of a naïve individual under the assumption of perfect knowledge of the environment led to an underestimation of the selection strength associated with a given environmental feature. Decomposing the redistribution kernel into its habitat- and movement-selection components helps with understanding differences between the omniscient and learner models (Fig. 4). Initially, the learner’s habitat-selection function is constant outside of the areas it has visited or that fall within its perceptual range due to lack of spatial knowledge. Its movement kernel is correspondingly more dispersed: the updated gamma distribution has a larger shape parameter than that of the omniscient model (shape_O_ = 0.62 vs shape_L_= 0.82), placing less probability mass immediately around the current location and increasing the median step length from 577 m to 846 m (Fig. 4A). As the individual accumulates spatial knowledge, the habitat-selection function in the learner model becomes more heterogeneous (Fig. 4B). Where knowledge is still absent, the habitat-selection function contributes a constant, and within known areas relative selection reaches more extreme values than in the omniscient model. The movement component is estimated once per model and therefore does not change between early and late steps.

**Figure 3.**
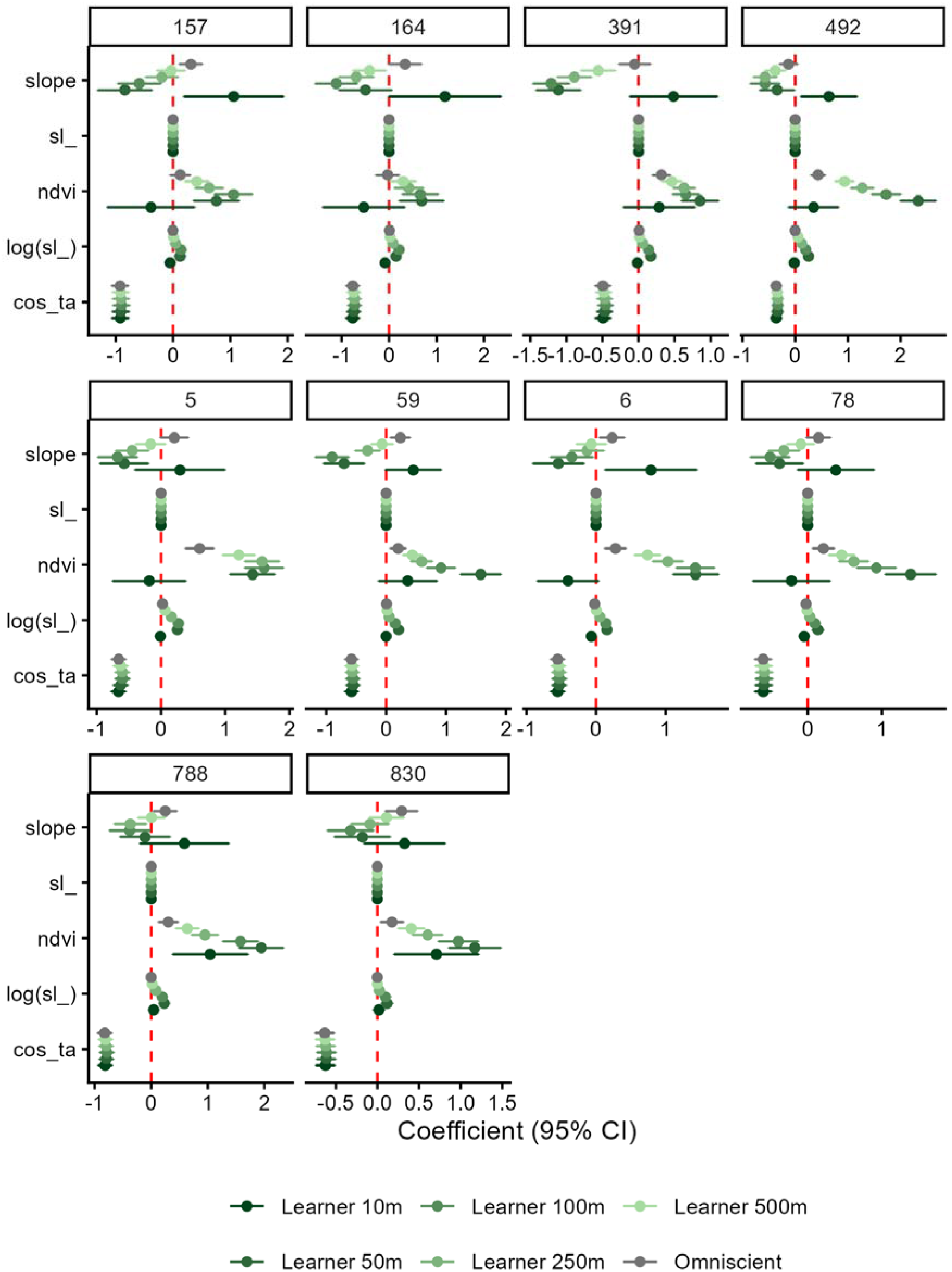
SSF coefficient estimates (±95% CI) for 10 juvenile red kites during dispersal from the natal home range. Covariates include step length, log step length, cosine of turn angle, NDVI, and slope. Models assume either omniscient knowledge of habitat covariates (grey) or learning-based, locally updated knowledge with perceptual radii from 10 to 500 m (green).

**Figure 4.**
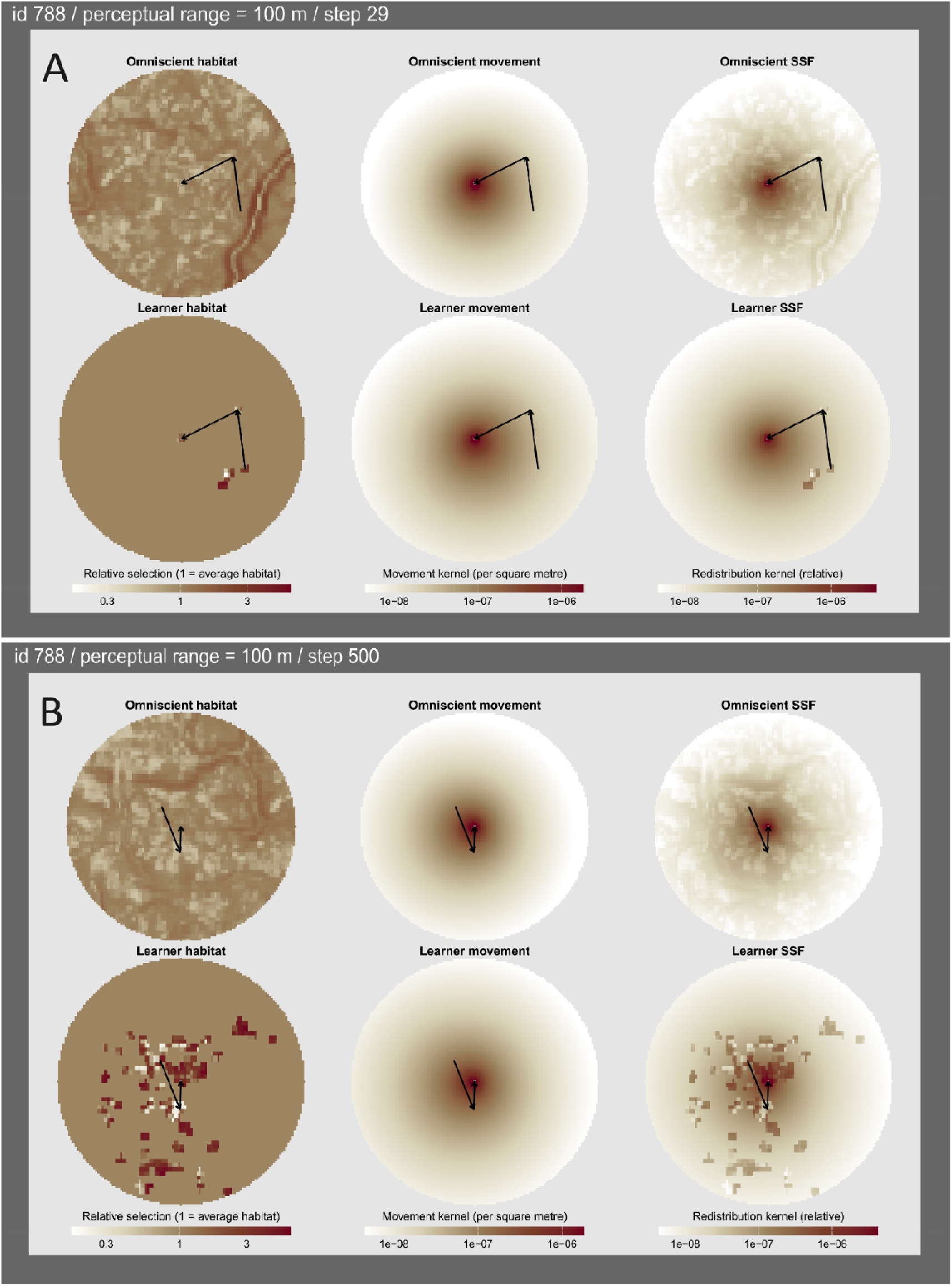
Representative model components (habitat-selection function, movement kernel, and their product) for a juvenile red kite with step-selection analyses assuming either an omniscient individual (top row) or a learner with a 100 m perceptual radius (bottom row). Panel A shows the three model components at an early stage of the trajectory (step 29), whereas Panel B shows the corresponding components at a later stage (step 500). Arrows indicate the two most recent steps of the trajectory. Colour scales are standardised within each kernel type across panels.

AIC identified an intermediate perceptual range as best supported in every individual. The learner model assuming a 100 m perceptual range had the lowest AIC in seven of ten birds and 50 m in the remaining three; the omniscient model was never best supported (Fig. 5). Support declined on both sides of the optimal perceptual range, mirroring the pattern found in the simulations, with median ΔAIC rising to 25 at 50 m, 42 at 250 m and 89 at 500 m. Notably, the smallest perceptual range tested (10 m, median ΔAIC 134) was supported no better than the omniscient model (median ΔAIC 122), indicating that a substantially underestimated perceptual range offers no advantage over ignoring learning altogether.

**Figure 5.**
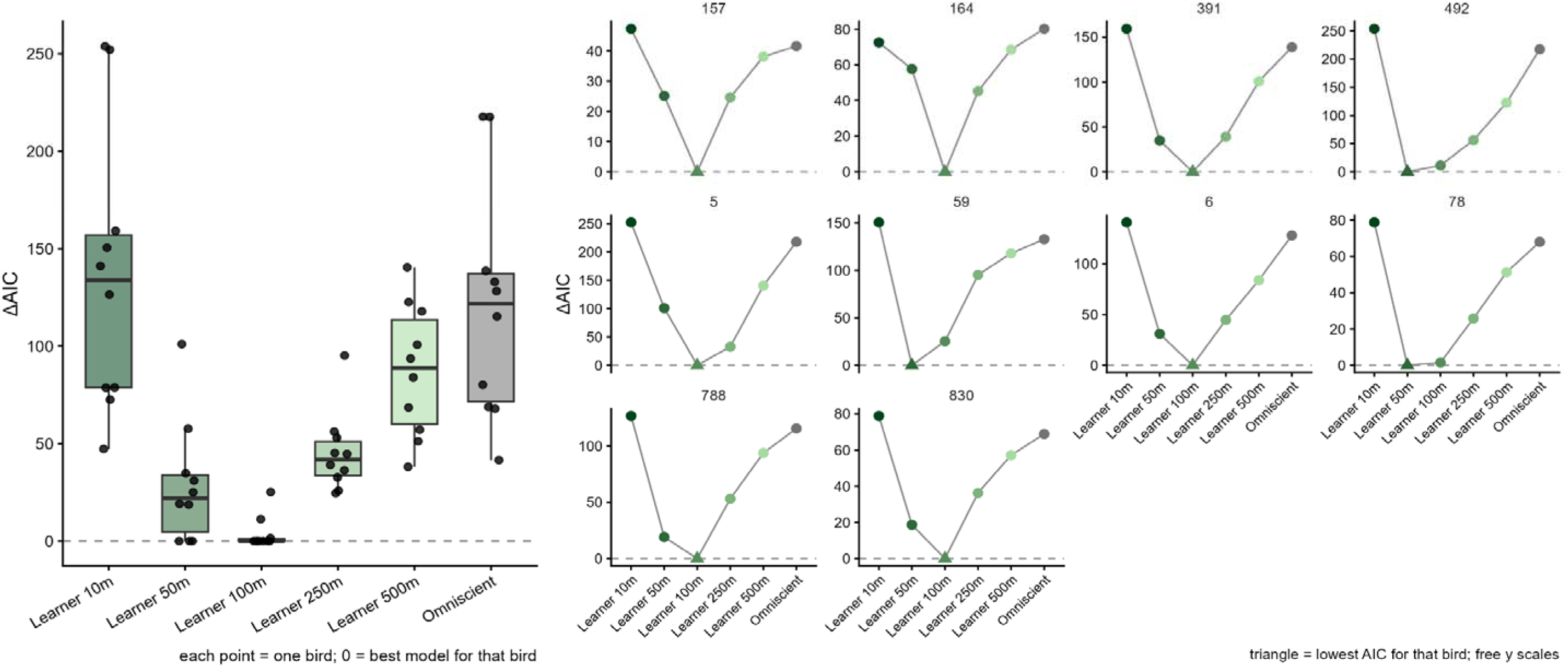
AIC-based comparison of learner and omniscient step-selection functions for ten juvenile red kites. For each individual, six step-selection functions were fitted to identical data: five learner models assuming perceptual ranges of 10, 50, 100, 250 and 500 m, and one omniscient model in which the entire landscape was known from the first step. ΔAIC is computed within individual, relative to the best-supported model for that bird, so that zero identifies the selected model. Left: distribution of ΔAIC across the ten individuals; boxes show the median and interquartile range, whiskers extend to 1.5× the interquartile range, and each point is one individual. Right: the same values shown separately for each individual (panel headings give bird identity), with the triangle marking the lowest-AIC model for that individual.

Cross-validation identified the same optimal perceptual range (Fig. S5). The learner SSF predicted held-out individuals better than the omniscient SSF across every perceptual range from 50 to 500 m, with the advantage being greatest at 100 m — the value also favoured by AIC. Mean differences in summed log-score (learner − omniscient) were negative (−7.43) at 10 m, 41.8 at 50 m, 53.2 at 100 m, 31.9 at 250 m and 13.1 at 500 m.

## 5 Discussion

Step-selection analysis (SSA) is a widely used approach for studying animal movement. In its standard formulation, environmental influences on movement decisions are inferred by comparing the characteristics of observed steps with those of a set of available alternatives (Thurfjell et al., 2014). However, this approach assumes that animals can perceive or know the environments represented by those observed and available steps. This assumption may be violated when individuals move through unfamiliar landscapes, such as after translocation or during natal dispersal, where decisions are shaped by both habitat preference and incomplete information. In such cases, standard SSA may attribute lack of movement to or toward highly attractive but unfamiliar locations as avoidance, biasing estimators of habitat-selection coefficients. Incorporating learning and information acquisition into movement analyses is therefore critical for understanding behaviour during post-release adjustment phases (Ebenhoch et al., 2019; Kemink & Kesler, 2013) and is becoming increasingly relevant as advances in tracking technology enable detailed study of juvenile dispersal and other early-life movements (Ponchon, 2026).

Here, we propose an extension of the step-selection framework in which environmental covariates are constructed in a way that is meant to reflect the animal’s learned landscape. In this approach, environmental features are considered known only when they fall within the area that the animal has previously experienced or could plausibly perceive. This distinction allows habitat selection to be estimated conditional on what the animal is likely to know at a given point in time. Importantly, the methodological advance proposed here does not require the development of a new modelling framework (Bracis et al., 2015; Fagan et al., 2023; Pollock et al., 2026; Ranc et al., 2022) or the addition of familiarity covariates (Oliveira-Santos et al., 2016). Rather, it represents a conceptual extension of standard SSA, where environmental covariates are dynamically updated to reflect the animal’s accumulated knowledge of the landscape. We also provide R code illustrating how this approach can be implemented within a standard SSA workflow.

Our simulations show that ignoring information constraints can lead to substantial underestimation of habitat-selection coefficients. The red kite case study lends further support to these results. Learner SSFs generally produced larger habitat-selection coefficients than omniscient models and consistently improved predictive performance relative to traditional SSFs. Orgeret et al., (2023) previously found that red kite habitat selection was weaker during the exploratory phase of dispersal than after red kites had established a territory. Our results offer a new interpretation of this pattern, suggesting that the noted weaker habitat preferences may have arisen from limited environmental knowledge, rather than from reduced selectivity itself.

Explicitly accounting for the learning process through which animals discover attractive environmental features (represented in our simulations by the “Easter egg”) also has important implications for the interpretation of movement patterns. Studies often infer attraction or avoidance of environmental features such as roads, anthropogenic food sources, or other landscape elements, implicitly assuming that individuals are already aware of their existence (Latham et al., 2011; Prokopenko et al., 2017; Reinking et al., 2019). While this assumption may be reasonable for resident animals that have had previous opportunities to experience and learn about local resources, it is less appropriate for juveniles, dispersers, or translocated individuals encountering a novel environment. For these animals, potentially important resources or risky areas must first be located before they can influence movement decisions. Ignoring this discovery phase may lead to biased estimators of habitat preferences and resource selection. In translocation contexts, ignoring how individuals learn about available resources may underestimate the importance of supplementary resources, as non-use may simply result from a lack of discovery rather than a lack of benefit (Greggor et al., 2024). Likewise, during dispersal, neglecting broad-scale learning processes may obscure how individuals gather information in unfamiliar environments and make settlement decisions (Clobert et al., 2009).

Beyond parameter estimation, learning also influenced movement behaviour. Learners used slightly larger areas, and their movements were less tortuous, with reduced revisitation and residence times compared with omniscient individuals, all consistent with increased exploratory behaviour while acquiring information about the landscape. In contrast, omniscient individuals were able to exploit favourable habitat immediately, resulting in more localised and sinuous movements. Notably, these differences were strongest in heterogeneous environments and largely disappeared in homogeneous landscapes, suggesting that the value of learning increases with environmental patchiness.

Decomposing the redistribution kernel into its habitat and movement components helped clarify where the two formulations diverge. Early in the trajectory, the omniscient model assigns habitat selection values across the entire accessible area, whereas the learner model does so only within areas that the individual previously visited or that fall within its perceptual range. As such, the learner’s early movement decisions are driven almost entirely by the movement kernel. As knowledge accumulates, the learner’s habitat-selection function progressively fills in, allowing it to have more influence on the individual’s movements. The movement kernel was also more dispersed in the learner model compared to the omniscient model. This result reflects how the two components jointly account for the same movements: a long step ending in distant but favourable habitat can be attributed to habitat selection only if the animal knows that habitat. In the learner model, those destinations are initially unknown, so habitat selection cannot explain large initial movements; however, the more dispersed movement kernel can accommodate them. The two components of the model (movement and habitat selection) are therefore not uniquely identified; what one component cannot account for, the other must. Ignoring learning therefore risks biasing not only habitat selection coefficients but also the selection-free movement parameters that are routinely extracted from step-selection functions to parameterise dispersal and connectivity models (Hofmann et al., 2023; Osipova et al., 2019).

It is interesting to consider the differential impacts of learning and memory on animal movements. Whereas learning requires more exploratory movements to learn about previously unfamiliar areas, memory allows animals to return to highly profitable locations that they previously visited. Ranc et al., (2022) found that memory promoted recursive movements and home-range formation. Incorporating learning in our SSFs led to larger habitat-selection coefficients, whereas incorporating memory or familiarity covariates often has the opposite effect (Oliveira-Santos et al., 2016; Ranc et al., 2022). This latter behaviour likely occurs because some of the spatial structure otherwise attributed to habitat preference is captured by the familiarity covariates, which promotes returns to previously visited areas. Our learner-based SSF approach differs in that we modify the habitat covariates to reflect what the animal knows, rather than introducing familiarity as an additional driver of movement. This allows habitat preferences to be estimated while accounting for the learning process that occurs as animals move through unfamiliar landscapes.

A central component of our framework is the definition of perceptual range. Our results suggest this parameter can be identified from the data rather than assumed: in the simulations AIC neatly recovered the generating perceptual range, and in the red kite application, AIC and cross-validation independently converged on the same value. Data-driven selection is therefore a practical default, complementary to *a priori* specification based on the biology of the study species, i.e. sensory abilities, movement mode, body size, and the structure of the environment (Fagan et al., 2017; Prevedello et al., 2011) which is best used to bound the set of candidate ranges rather than to fix a single value. Plausible ranges must also be considered relative to the scale of observed movement. When the perceptual range is much smaller than typical step lengths, both used and available locations will frequently fall outside the perceived area, leaving little contrast in the habitat covariates, although this constraint relaxes as the animal accumulates knowledge. Perceptual range is also unlikely to be the same for all environmental covariates. Some features, such as forest edges, roads, rivers, cliffs, or broad habitat types, may be detectable from relatively long distances (Olden et al., 2004). Others, such as fine-scale vegetation composition, prey abundance, or predation risk, may only be perceived at short range or after direct experience. Future applications of this framework could therefore allow different perceptual ranges for different covariates. This would better reflect the fact that animals may learn about different aspects of the environment at different spatial scales.

The effective perceptual range of an animal may also differ among individuals or change over time in response to external factors such as time of day or social information. In our framework, perceptual range determines how quickly and over what spatial extent environmental knowledge is acquired as the animal moves. It could therefore be treated as a tuneable parameter rather than a fixed constant, for instance estimated separately for each individual to capture inter-individual variation, or allowed to vary within an individual’s trajectory according to time of day (e.g. a smaller perceptual range at night than during the day for diurnal species). Beyond direct perception, conspecifics, heterospecifics, scent marks or vocalisations may provide cues about resources, risk, competitors, or suitable habitat (Jesmer et al., 2018; Loonstra et al., 2023). In this sense, social cues could extend the animal’s informational landscape beyond the area directly visited or perceived. This consideration may be particularly relevant for social species, territorial species, or translocated animals released near resident individuals (Mihoub et al., 2011). Incorporating social information into learning-based step-selection models could therefore improve inference where movement decisions depend strongly on public information or conspecific attraction.

Another promising extension of this approach would be to incorporate experience on top of the learning process. Doing so would help address the question of whether learning should be represented solely as knowledge of previously visited locations, or whether animals also acquire more general information about the environmental conditions they encounter. Our current framework assumes that individuals gradually build a spatial representation of the landscape as they move. However, learning may also allow animals to associate particular environmental conditions with positive or negative outcomes and to apply this information in novel contexts. For example, a prey animal that experiences a predator encounter in dense shrub cover may learn not only that a specific location is dangerous, but also that dense vegetation is generally associated with higher predation risk. Our framework also currently assumes that acquired spatial knowledge remains valid indefinitely, whereas real landscapes change. Thus, we might expect the value of information gained from an individual’s past movements to decrease over time (Spencer, 2012). Capturing these processes—the ability to infer the general value of different landscape features across space and information decay over time—would be valuable but difficult to do within the SSF/conditional logistic regression modelling framework. One of the main benefits of our learning model is its accessibility – models can be fit using standard statistical software and using tools that ecologists are already familiar with.

Animals do not always make decisions with complete knowledge of their surroundings, and by incorporating learning into SSA, we provide a simple approach to distinguish habitat preferences from limitations in environmental knowledge. Such a framework may prove especially valuable whenever animals must make decisions under uncertainty, from dispersal and migration to biological invasions, translocations, and adaptation to rapidly changing environments. We hope this work provides a foundation for future efforts towards building a more mechanistic understanding of animal movement.

## Acknowledgements

This work was supported by the Swiss National Science Foundation (grants 310030_212469 and 31003A_169668 to MUG). JF was supported by the National Aeronautics and Space Administration (award 80NSSC21K1182) and received partial salary support from the Minnesota Agricultural Experiment Station. Experimental licences for GPS tagging of red kites were provided by the Amt für Lebensmittelsicherheit und Veterinärwesen (LSVW) of the Canton of Fribourg (permit no. 2017_29_FR) and the Federal Office for the Environment (FOEN). We thank Patrick Scheler, Urs G. Kormann, Valentijn van Bergen and all the many field assistants and students who contributed to the collection and curation of the red kite GPS tracking dataset.

