## Supplementary figures 1-5 for "Integrating learning in movement using step-selection analyses"

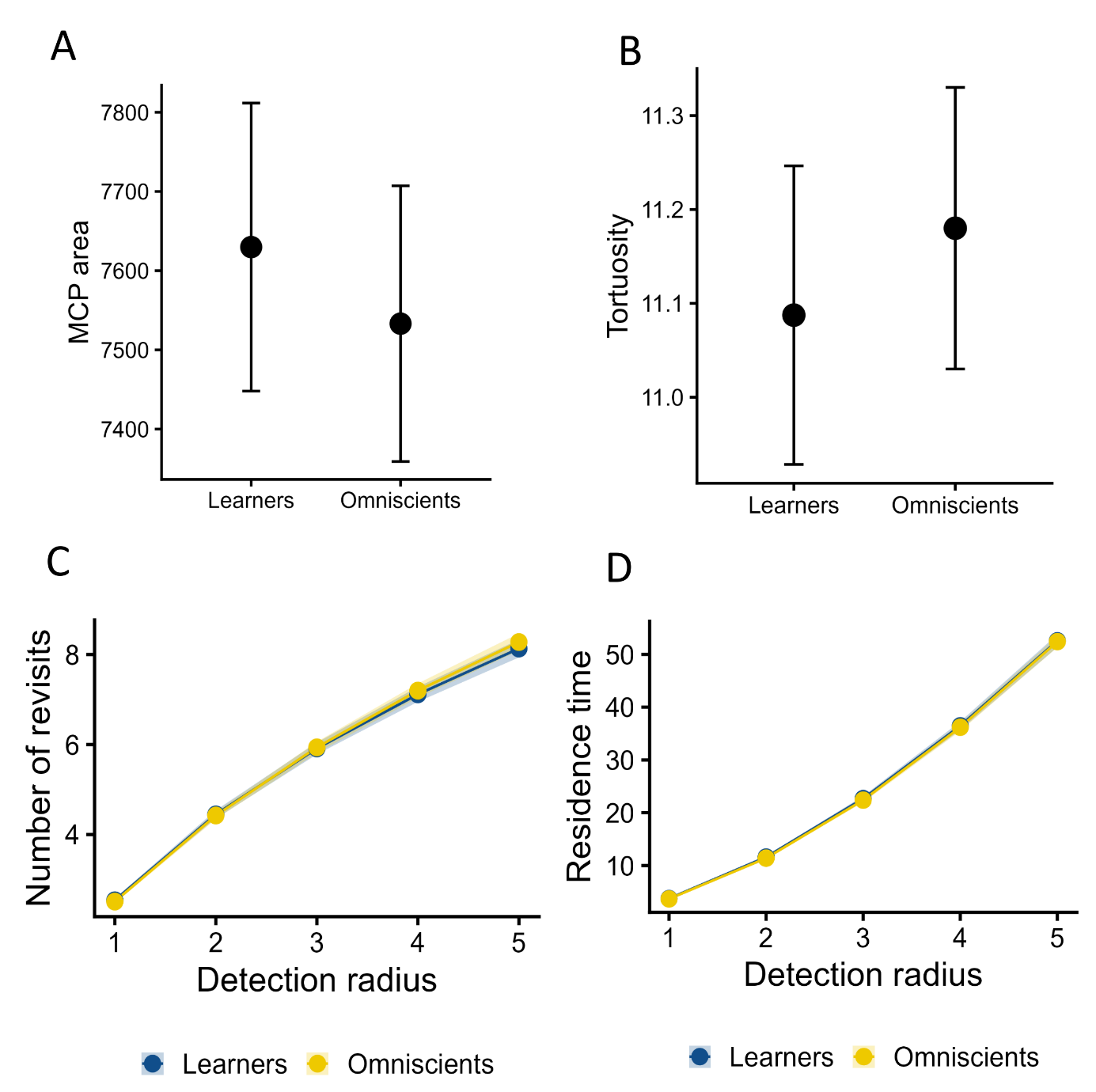

Figure S1 Movement metrics from 500 simulated trajectories under omniscient and learner movement strategies in a homogeneous landscape. Trajectories were simulated with positive selection for favourable habitat (β_habitat_ = 0.5) and identical movement parameters in both scenarios. In the traditional "omniscient" scenario, individuals evaluate environmental conditions at every available step. In the "learner" scenario, individuals have only partial knowledge: a candidate step is either known (previously visited, or within the perceptual range of a previously visited location) or unknown, in which case it takes the naive prior. (A) MCP area. (B) Path tortuosity, the ratio of path length to straight-line displacement over a moving window of 100 consecutive locations, averaged across windows within an individual. (C, D) Movement recursiveness across increasing spatial scales, quantified as the number of revisits (C) and residence time (D). Points are means across the 500 individuals per strategy and error bars are 95% confidence intervals for those means.

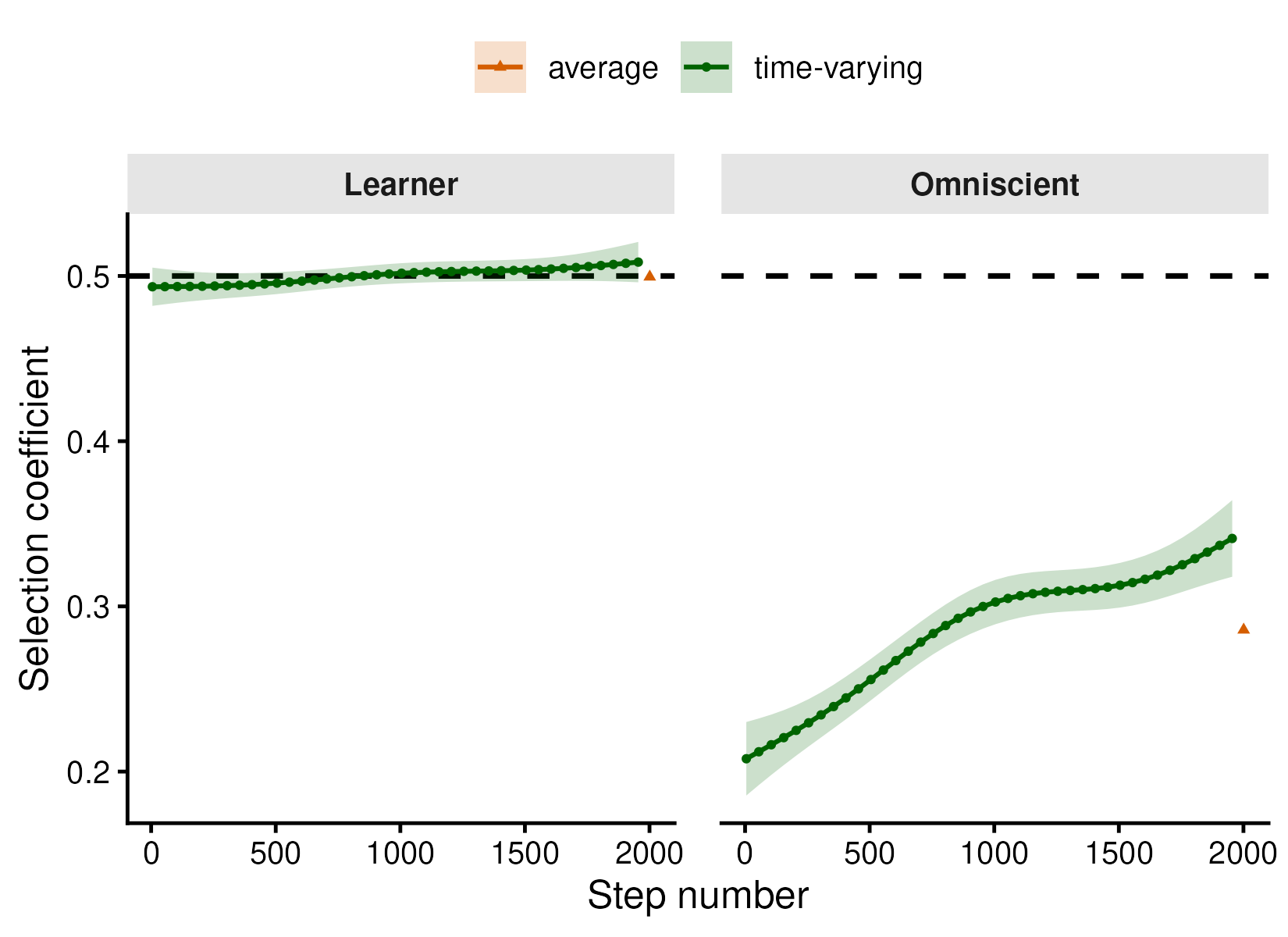

Figure S2 Recovery of habitat selection coefficients from learner-simulated trajectories under different SSF formulations.	We assessed how accurately the true habitat selection coefficient (β = 0.5) used to simulate learner movement trajectories is recovered using two analytical approaches. The left panel shows results from a step-selection function (SSF) that explicitly incorporates learning, whereas the right panel shows results from a traditional “omniscient” SSF assuming perfect environmental knowledge. Green points represent time-varying coefficient estimates averaged across 500 simulations, obtained from GAMs at successive steps along the trajectory. Orange triangles show the corresponding constant (time-invariant) estimates, also averaged across simulations. Shaded bands indicate 95% Monte Carlo interval for that mean. Horizontal dashed lines denote the true coefficient values used in the simulations.

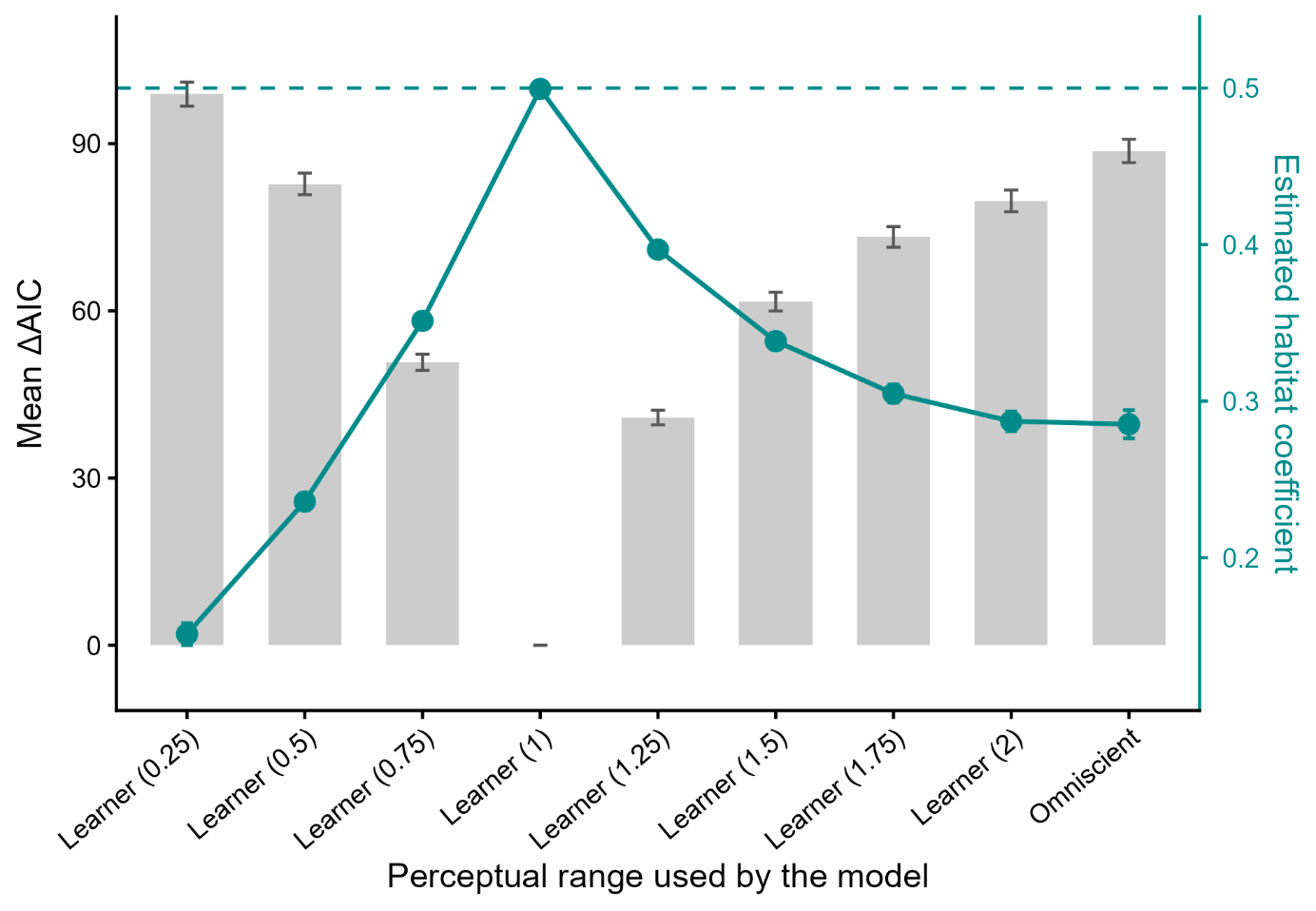

Figure S3 Model selection and coefficient recovery across candidate perceptual ranges. Learner step-selection functions were fitted at eight perceptual ranges (0.25–2 map units) and compared with an omniscient model in which the habitat covariate was known everywhere. Within each simulated individual all nine models were fitted to identical strata and carry the same number of parameters, so ΔAIC equals twice the difference in partial log-likelihood. Grey bars show the mean ΔAIC across 500 individuals, relative to the best-supported model for that individual (0 = best; lower is better). Coloured points and line show the mean estimated habitat-selection coefficient, read against the right-hand axis; the dashed horizontal line marks the true value used to generate the trajectories (β = 0.5). Error bars are 95% Monte Carlo intervals for the mean across individuals. The black tick on the x axis marks the perceptual range at which the trajectories were simulated (1 map unit).

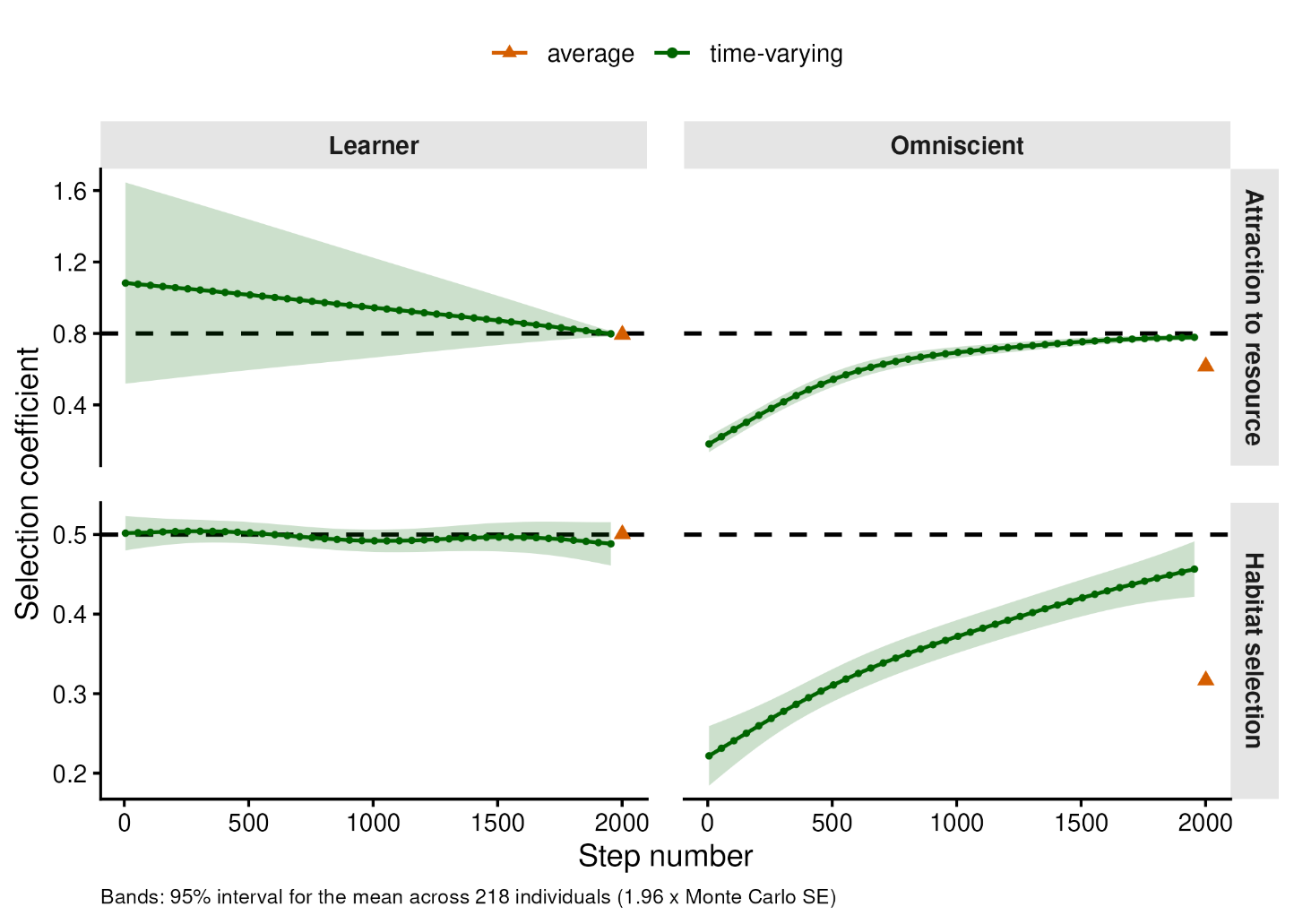

Figure S4 We assessed how accurately the true habitat-selection coefficient (β = 0.5) and coefficient associated with the relative angle to the source (i.e. the Easter egg; β = 0.8) are recovered using two analytical approaches. Columns show results from a learning-informed SSF (left) and a traditional “omniscient” SSF assuming perfect environmental knowledge (right), while rows correspond to selection for the continuous habitat covariate (bottom) and for the relative angle to the source (top). Green points represent time-varying coefficient estimates averaged across 500 simulations, obtained from GAMs at successive steps along the trajectory. Orange triangles show the corresponding constant (time-invariant) estimates, also averaged across simulations. Shaded bands indicate 95% Monte Carlo interval for that mean. Horizontal dashed lines denote the true coefficient values used in the simulations.

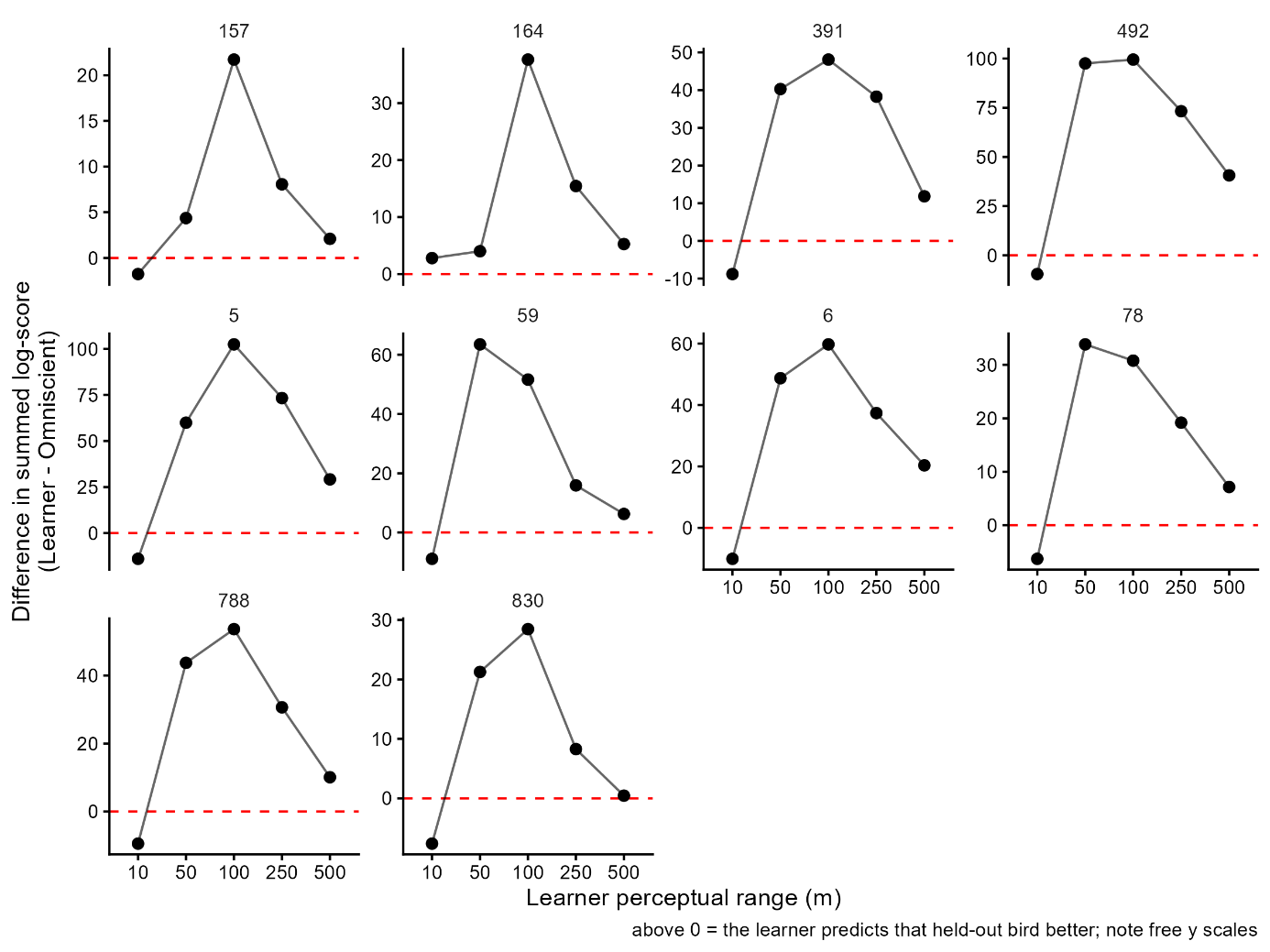

**Figure S5** Leave-one-individual-out cross-validation comparing the out-of-sample predictive performance of learner and omniscient SSA models across a range of assumed perceptual ranges, including unrealistically small buffer sizes (10–500 m). For each individual red kite (panel), the model was fitted to data from all other individuals and used to predict the held-out individual's steps; the y-axis shows the mean difference in summed log-score between the two models (learner − omniscient) at each buffer size, with values above the dashed red line indicating better predictive performance for the learner model. Predictive performance was consistently poor at the smallest, biologically unrealistic buffer (10 m), improved sharply as buffer size increased, and peaked at intermediate values (50–100 m) for most individuals.
